# Modeling Parkinsonian Freezing and Deep Brain Stimulation Effects in a Basal Ganglia Network

**DOI:** 10.64898/2026.07.21.738458

**Authors:** Nahal Habibizadeh, Madeleine Bartlett

**Affiliations:** Cheriton School of Computer Science, University of Waterloo; National Research Council Canada

**Keywords:** Parkinson’s disease, basal ganglia, action selection, freezing of movement, dynamic neural fields, dopamine depletion, thalamic gating, deep brain stimulation, computational neuroscience

## Abstract

The precise mechanism through which Deep Brain Stimulation (DBS) mitigates freezing episodes in Parkinson’s disease remains unknown. We modelled Parkinsonian freezing using a model of the basal ganglia that adopts a Dynamic Neural Field network for simultaneous action selection (selecting *which* action is executed) and specification (resolving the continuous parameters governing *how* that action is performed). Action selection success was based on whether activation surpassed a threshold. The model robustly differentiated healthy and dopamine-depleted (Parkinsonian) conditions, producing stable action selection under healthy dopamine and impaired, freeze-prone dynamics following depletion. We then modelled DBS as a scaled reduction in afferent drive to the subthalamic nucleus and globus pallidus internus. While this formulation has succeeded in previous computational work, it did not yield consistent restoration of action selection within our framework, motivating future investigation of alternative DBS formulations.

## Introduction

Parkinson’s disease (PD) is a progressive neurological disorder characterized by a complex constellation of motor and cognitive symptoms, one of which is a freezing phenomenon affecting both gait and speech (Erro et al., 2014; Vercruysse et al., 2014). The primary neuropathological mechanism underlying PD is the degeneration of dopaminergic neurons in the substantia nigra pars compacta (SNc), leading to reduced dopamine levels in the striatum which disrupts the function of the basal ganglia (BG) (Goetz & Pal, 2014). The BG is widely understood as an action selection and gating mechanism that resolves competition between parallel motor and cognitive programs by facilitating a single context-appropriate action selection while suppressing alternatives (Mink, 1996; Redgrave et al., 1999). This selection process is implemented through interactions among the basal ganglia nuclei, including the striatum, subthalamic nucleus (STN), globus pallidus externus (GPe), and globus pallidus internus (GPi), which together shape inhibitory and excitatory signaling to regulate action gating. Dopaminergic modulation within the striatum plays a critical role in this process via two major pathways: the direct pathway which facilitates action selection and is modulated by D1 receptor–expressing neurons, and the indirect pathway which regulates competition via the STN-Gpe control loop and is modulated by D2 receptor–expressing neurons (Obeso et al., 2008). Dopamine depletion compromises the action gating process, resulting in pathological competition between candidate actions and, in severe cases, failures of action initiation that manifest as freezing. Freezing phenomena provide a particularly informative behavioral window into basal ganglia dysfunction, as they reflect a failure of action initiation rather than an incorrect action choice (Ackermann et al., 1997). In the clinical realm, Deep Brain Stimulation (DBS) has emerged as an effective intervention to alleviate many motor symptoms of PD, particularly when applied to STN or GPi, regions that exhibit abnormal activity in PD patients (Liu et al., 2014). However, despite its widespread clinical use and proven successes, the precise mechanisms by which DBS restores function remain the subject of ongoing experimental and computational investigation (Ashkan et al., 2017; Chiken & Nambu, 2016).

A wide range of computational models have been proposed to study the effects of DBS, spanning biophysical, patientspecific, and functional levels of description (Agnesi et al., 2013; Herrington et al., 2016; McIntyre & Foutz, 2013; Senft et al., 2018). Biophysical models explicitly simulate stimulation-induced electric fields and neural activation, demonstrating that DBS can differentially suppress somatic firing while activating axonal pathways in regions such as the STN and GPi (McIntyre et al., 2004; Miocinovic et al., 2006). While these approaches provide critical mechanistic insight, they require detailed anatomical and electrical modeling and are not designed to address basal ganglia computation or action selection. At a functional level, circuit-based models abstract away from pulse-level stimulation and instead characterize DBS as a modulation of effective connectivity within basal ganglia circuits. Functional level computational models of the basal ganglia–thalamocortical loop have successfully linked lowered dopamine conditions to freezing by demonstrating degraded action selection in Parkinsonian models (Gurney et al., 2001b; Senft et al., 2018). In these models, the selection process is implemented through tightly coupled excitatory, inhibitory, and disinhibitory pathways spanning the basal ganglia. Within this framework, cortical representations of candidate actions compete at the striatal level, while downstream basal ganglia output nuclei regulate thalamic disinhibition to permit action selection (Chiken & Nambu, 2016; Gurney et al., 2001a). In this study, we adopt the formulation proposed by Senft et al. (2018), which models DBS as a scaled reduction in afferent drive to the STN and GPi. This approach is grounded in the hypothesis that therapeutic DBS suppresses pathological basal ganglia input to the thalamus and restores proper action gating, providing a tractable framework for observing DBS effects within a BG model(Ashkan et al., 2017).

In this work, we extend recent advances in computational accounts by using a novel instantiation of a Dynamic Neural Field basal ganglia (DNF-BG) model proposed by Bartlett et al. (2025). In contrast to traditional models that focus primarily on discrete action selection, this framework characterizes the basal ganglia as a collection of discrete action channels whose internal dynamics specify how selected actions are carried out. Inputs to the model consist of distributions of salience over continuous action parameter spaces (e.g. speed), with one distribution per action channel (Saha et al., 2026). Leveraging a one-channel version of this architecture, we explicitly model Parkinsonian freezing as a failure of thalamic action gating to identify any point in the action parameter space as having sufficient salience for execution. By coupling the DNF-BG model to an explicit thalamic gating mechanism, we compare action selection performance under healthy and dopamine-depleted conditions while directly quantifying selection failures associated with freezing. This formulation enables robust differentiation between healthy and Parkinsonian regimes. Building on this validated distinction, we also conducted an exploratory investigation of DBS, examining whether the DBS formulation established by Senft et al. (2018) can alleviate freezing symptoms under dopamine-depleted conditions.

## Methods

### Basal Ganglia Model

As mentioned above, all experiments adopt the DNF-BG model developed by Bartlett et al. (2025). The model represents continuous action spaces using Spatial Semantic Pointers (SSPs; Voelker et al., 2021) and includes anatomically grounded neuron populations corresponding to striatum (D1 and D2), STN, GPe, and GPi (see Figure 1). Rather than treating action selection purely as competition between discrete actions, the DNF–BG model organizes the basal ganglia into parallel action channels whose internal dynamics determine the continuous parameters governing *how* an action is expressed. As such, potential actions are represented as continuous salience distributions over the action parameter space, where higher salience values reflect stronger support for particular action realizations. Within this architecture, competition between channels and refinement of action parameters are jointly resolved through DNF dynamics, producing sharpened peaks around the most strongly supported action saliences (Saha et al., 2026; Schöner & Spencer, 2016).

**Figure 1.**
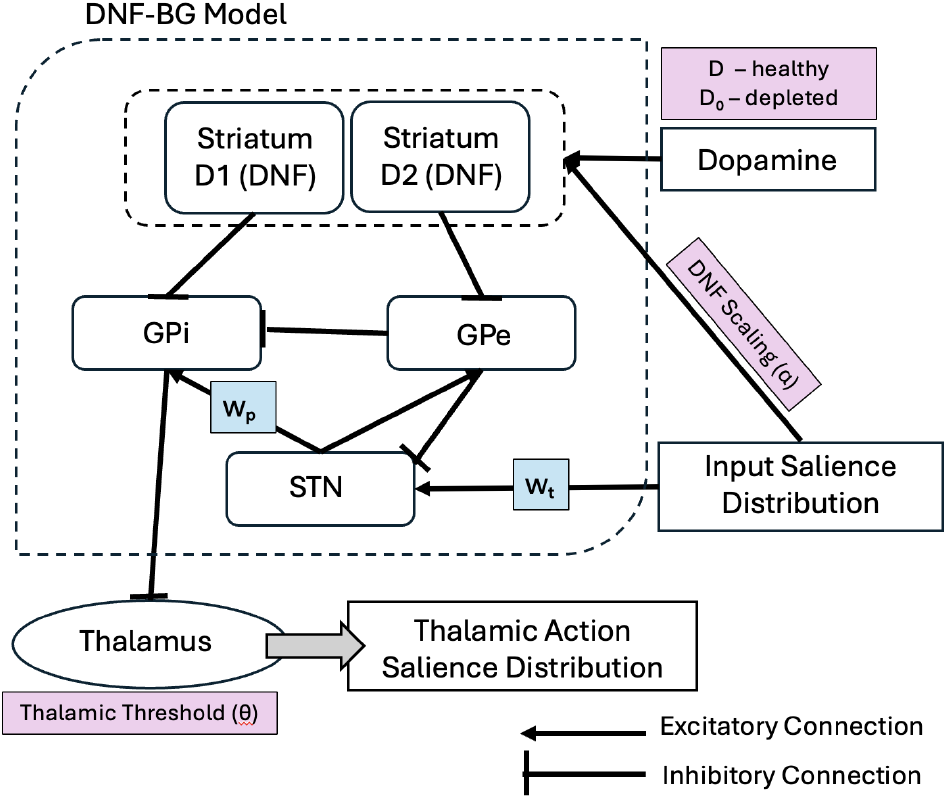
Schematic of the DNF-based basal ganglia– thalamus action selection model. A salience input distribution is scaled by the DNF gain *α* and, together with a dopamine signal (*D* for healthy, *D*_0_ for depleted), drives striatal D1/D2 dynamic neural fields (DNFs). The same input provides excitatory drive to the subthalamic nucleus (STN), scaled by *w*_*t*_, while STN excitation of GPi is scaled by *w* _*p*_ in the DBS formulation. GPi inhibits thalamic activity, where mutual inhibition and a threshold *θ* determine action selection

For the purpose of our studies, we introduced an (*α*) parameter that globally scales the strength of recurrent DNF dynamics in the striatum. The value of *α* determines how strongly intrinsic striatal dynamics contribute to action selection across dopamine levels, with higher values increasingly supporting selection even under dopamine depletion and potentially obscuring freezing-like behavior.

### Thalamus

In our adaptation of the DNF-BG, action selection is completed through an explicit thalamic gating mechanism implemented using Nengo’s built-in `nengo.networks.Thalamus` network (Bekolay et al., 2014). This network performs action selection through mutual inhibition: each dimension of the network corresponds to a candidate action and receives inhibitory input from the basal ganglia’s output, with stronger inhibition reflecting weaker support for a given action. The continuous salience distribution encoded in the BG’s output is passed into this network. The thalamic network consists of a set of channels that take, as input, the decoded salience values for a finite set of equally spaced points across the continuous action space. Each channel’s thalamic activity is disinhibited in proportion to its corresponding action’s relative salience, such that the most strongly supported point in the action space emerges as the least inhibited channel. A channel is considered selected if, at the end of the simulation window, its activity exceeds a decision threshold (*θ*). Then, the point in the action space corresponding to this channel is taken as the selected action. If the decision threshold is not exceeded by any channel, we consider it as a failure of action selection and a freezing occurrence. To directly compare healthy and freeze-prone pathologies in the same model, we require a model such that, under dopamine-depleted conditions, the thalamic network will fail to successfully select an action. Therefore, the thalamic threshold (*θ*) determines whether a given dopamine level is sufficient for successful action selection. A schematic of the full basal ganglia–thalamocortical architecture is shown in Figure 1.

### Dopamine Modulation

Dopamine is implemented as a global gain parameter that modulates relative contributions of the direct and indirect pathways by scaling activity in the D1 and D2 populations, as shown in Figure 1. High dopamine values increase effective signal-to-noise and support decisive winner-take-all selection, whereas reduced dopamine flattens salience representations and destabilizes competition (Bartlett et al., 2025; Humphries et al., 2012). To achieve a successful freezing model with the DNF-BG, we assess whether the framework can distinguish healthy and Parkinsonian dynamics under conditions that are otherwise identical, differing only in dopamine level. To distinguish dopamine regimes, we introduce fields *D* (healthy) and *D*_0_ (Parkinsonian) which correspond to high and low dopamine ranges used to emulate each condition, respectively.

### Action Selection Success

Based on the definition of success of action selection introduced above, we quantify action selection using a selection count metric defined as the number of trials (out of 25) in which an action was selected within the simulation time window of one second. This metric is computed separately for healthy dopamine conditions (Healthy_selected_), dopamine-depleted Parkinsonian conditions (PD_selected_), and dopamine-depleted conditions with DBS applied (DBS_selected_).

All simulation code, parameter sweep configurations, and result data are available at https://github.com/nahalhz/nhabibiz_cogsci_2026.

## Study 1: Parkinsonian Freezing Model

### Method

The goal of this study was to model freezing within the DNFBG model by establishing a model that exhibited substantial selection failures under dopamine-depleted conditions. We conducted a grid search over the (*α*) and (*θ*) parameters, testing the selection rate over 6 × 6 combinations of high vs. low dopamine. The goal of the grid search was to identify the optimal configuration of thalamic threshold *θ* and DNF scaling factor *α* that maximized the difference in action selection counts between a range of healthy (high dopamine) and Parkinsonian (dopamine-depleted) dopamine levels. In total, we searched an *α×θ*×*D*×*D*_0_ grid space, testing 432 total configurations (see Table 1 for parameter values included in the search). We refer to each parameter configuration in the grid search as an experiment. Each experiment consisted of 25 trials per dopamine condition, where each trial was a single action-selection attempt. Each trial produced a thalamic output of a 1 ×400 vector of action salience values. From there, we were able to conclude the success or failure of a trial’s action selection based on the method described in the *Thalamus* section. Finally, for each dopamine condition, a selection ratio (*R*) is defined to compare across experiments to find the configuration where the differentiation between healthy and Parkinsonian conditions is most substantial. The larger the ratio *R* of an experiment, the more successful the configuration.

**Table 1:** Parameter ranges used in the model grid search. Shaded rows indicate parameters varied in Study 2 (DBS), while unshaded rows indicate parameters varied in Study 1.

| Parameter | Values |
| --- | --- |
| $D$ | {1.0, 1.1, 1.2, 1.3, 1.4, 1.5} |
| $D_0$ | {0.08, 0.09, 0.10, 0.11, 0.12, 0.13} |
| $\theta$ | {0.2, 0.3, 0.4, 0.5} |
| $\alpha$ | {0.7, 0.8, 0.9} |
| $w_t$ | {0.5, 0.6, 0.7, 0.8, 0.9} |
| $w_p$ | {0.5, 0.6, 0.7, 0.8, 0.9} |

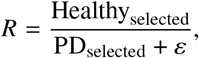

where ε is a small constant added to avoid division by zero.

### Results

The grid search revealed that thalamic thresholds in the range *θ* ∈ [0.4, 0.5] DNF scaling factors *α* ∈ [0.8, 0.9] consistently yielded high healthy selection counts alongside suppressed Parkinsonian selection as demonstrated by their high selection ratio value *R* (see Figure 2). Within the high-separation region of the parameter space in Figure 2, we collapsed results across all healthy *D* and dopamine-depleted *D*_0_ conditions in order to examine variability. Figure 3 shows the mean number of actions selected under healthy and dopamine-depleted conditions, with box-plots depicting the spread of action selection counts across experiments. These results demonstrate that these configurations exhibit stable selection dynamics. For the following DBS study, we selected a representative parameter configuration from among those with the highest *R*: *θ* = 0.4, *α* = 0.9, healthy dopamine *D* = 1.2, and dopamine-depleted *D*_0_ = 0.10. Statistical analyses to validate differences between the representative healthy and Parkinsonian conditions were conducted as part of Study 2, leveraging the repeated iterations performed using the same configuration.

**Figure 2.**
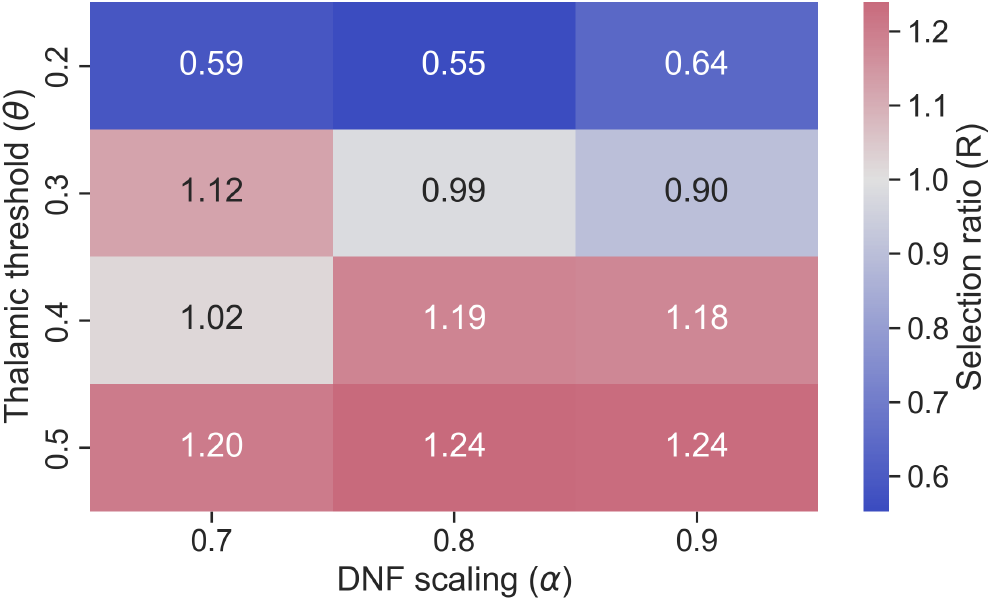
Healthy-to-Parkinsonian action selection ratio (*R*) across thalamic threshold (*θ*) and DNF scaling factor (*α*). Higher values indicate stronger differentiation between healthy and Parkinsonian dopamine levels, collapsed across *D* and *D*_0_.

**Figure 3.**
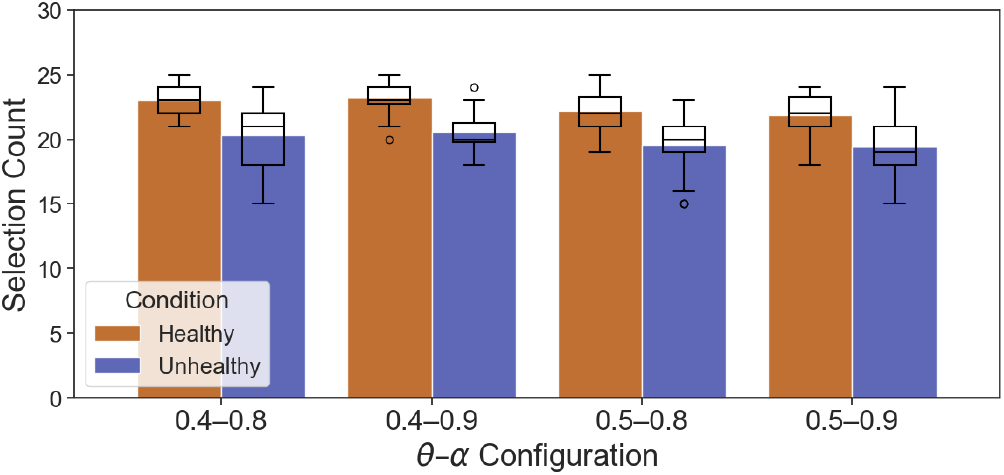
Action selection performance for the four highest-scoring *θ*–*α* combinations identified from the grid search. Bars indicate the mean number of selected actions (out of 25) under healthy (*D*) and dopamine-depleted (*D*_0_) conditions, while overlaid box-plots illustrate experiment-level variability across dopamine ranges per condition.

## Study 2: Deep Brain Stimulation Exploratory Study

### Method

To explore a proposed mechanism through which DBS alleviates freezing within the DNF-BG framework, we adopted an established modeling approach based on Senft et al. (2018) which parameterizes DBS as a modulation of activity within basal ganglia nuclei rather than explicitly modeling the delivery of stimulation current. Specifically, the stimulation is parameterized by the scaling factors *w*_*t*_ and *w* _*p*_ (see Figure 1). Reducing *w*_*t*_ decreases excitatory drive to the STN, while reducing *w* _*p*_ weakens inhibitory output from the GPi to the thalamus. This formulation reflects the hypothesis that DBS suppresses pathological basal ganglia activity affecting thalamic gating, restoring successful action selection. Treating DBS as a controlled modulation of information flow through the basal ganglia–thalamocortical loop enables systematic exploration of its effects without committing to a specific biophysical mechanism, which varies between patients in clinical settings (Ashkan et al., 2017; McIntyre & Foutz, 2013).

Following this framework, the baseline values (*w*_*t*_ = 1.0, *w* _*p*_ = 0.9) correspond to the Parkinsonian condition in the study, and DBS is simulated by reducing these parameters, thereby, scaling down the afferent signals (*w*_*t*_ < 1.0, *w* _*p*_ < 0.9). In the basal ganglia–thalamocortical model of Senft et al. (2018), the parameter settings *w*_*t*_ = 0.8 and *w* _*p*_ = 0.6 were reported as the most effective to compensate for dopamine-depleted conditions. Rather than adopting these values directly, we employ a more cautious approach by systematically exploring the parameter space thoroughly in our study.

#### Grid search Design

For the DBS condition, we performed a grid search over afferent scaling to the STN *w*_*t*_∈ [0.5, 0.9] and GPi *w* _*p*_ ∈ [0.5, 0.9] both varied in increments of 0.1 (see Table 1). For every (*w*_*t*_, *w*_*p*_) parameter pair, each experiment consisted of three conditions (healthy, Parkinsonian and DBS), with 25 trials per condition. To enable statistical evaluation of DBS effects, each experiment was repeated *n* = 20 times under identical parameter configurations for the entire grid search. This design enabled robust independent comparison of healthy, Parkinsonian, and DBS-modulated dynamics under matched conditions and supported significance testing of observed differences.

Using the representative parameter configuration from Study 1 (*θ* = 0.4, *α* = 0.9, *D* = 1.2, *D*_0_ = 0.10) across all experiments, DBS was applied to the Parkinsonian model and evaluated by examining changes in action selection relative to the untreated Parkinsonian condition. For each repeated experiment, we recorded whether the DBS condition produced a higher action selection count than the corresponding Parkinsonian condition. The proportion of repeated experiments satisfying this criterion was then used to summarize DBS performance across the parameter search:

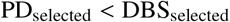

### Results

With the experimental design of the DBS grid search allowing for repeated simulations, we were able not only to assess the statistical significance of DBS-related effects, but also to further validate the healthy and Parkinsonian model configuration identified in Study 1. Non-parametric statistical analysis across repeated simulations (*n* = 20) using a Mann–Whitney *U* test revealed a highly significant difference in action selection counts between healthy and Parkinsonian conditions (*U* = 373.0, _*p*_ = 1.11 × 10^-6^), accompanied by a large effect size (*r*_*rb*_ = 0.865). Together, these findings validate the DNF–BG model as a robust and consistent computational account of dopamine-dependent action selection and freezing dynamics over continuous action spaces.

Figure 4 summarizes the percentage of successful DBS experiments across the (*w*_*t*_, *w* _*p*_) configurations (*n* = 20), based on the definition of success above, indicating reduced action selection failures compared to the Parkinsonian model. The best-performing configuration was *w*_*t*_ = 0.60 and *w* _*p*_ = 0.70, with improvement observed in 12 of 20 experiments (60%). However, we did not identify any region of the parameter space in which DBS reliably reduced freezing rates, as observed effects varied substantially across parameter configurations.

**Figure 4.**
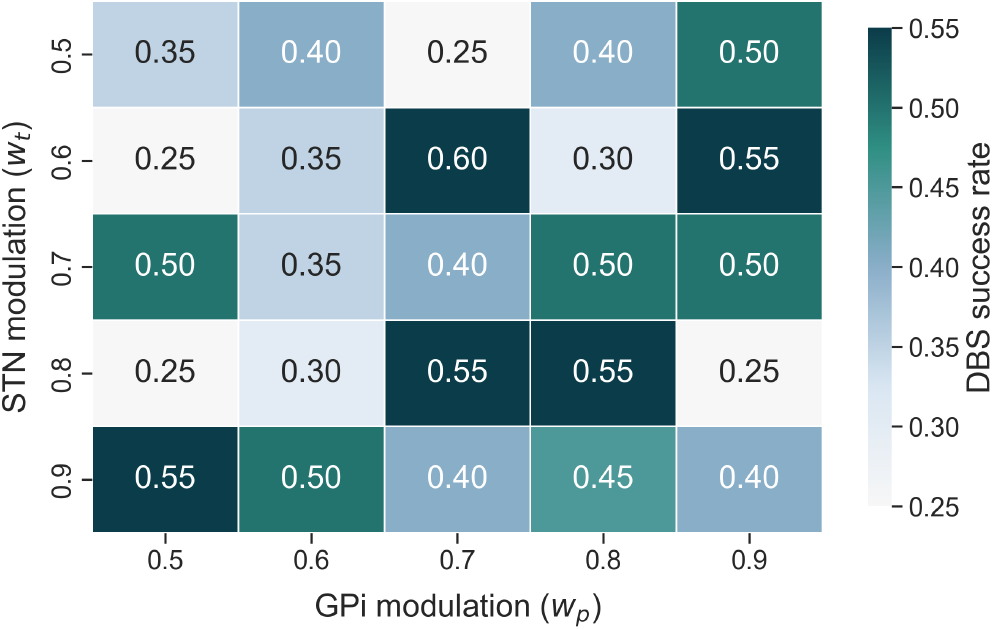
DBS success rate across GPi (*w* _*p*_) and STN (*w*_*t*_) modulation parameters. Values indicate the fraction of trials (out of 20) in which DBS selection performance exceeded the dopamine-depleted condition.

We conducted a closer examination of the effect of DBS on action selection for the best-performing configuration (*w* _*p*_ = 0.70, *w*_*t*_ = 0.60) across repeated runs. Figure 5 shows that the DBS condition produced only slight improvement of mean action selection counts relative to the dopamine-depleted condition (DBS: (*µ* = 20.70, *IQR* = 2.25) vs. Parkinsonian: (*µ* = 20.60, *IQR* = 3.00); *n* = 20), and both conditions remained substantially below healthy performance (Healthy: (*µ* = 23.25, *IQR* = 1.25). As such, action selection under the DBS condition remained statistically indistinguishable from the dopamine-depleted condition, as confirmed by a Mann–Whitney *U* test (*U* = 213.5, _*p*_ = 0.36, *r*_rb_ = 0.06). Together, these results indicate that within the explored parameter space, DBS did not robustly restore healthy-like action gating dynamics.

**Figure 5.**
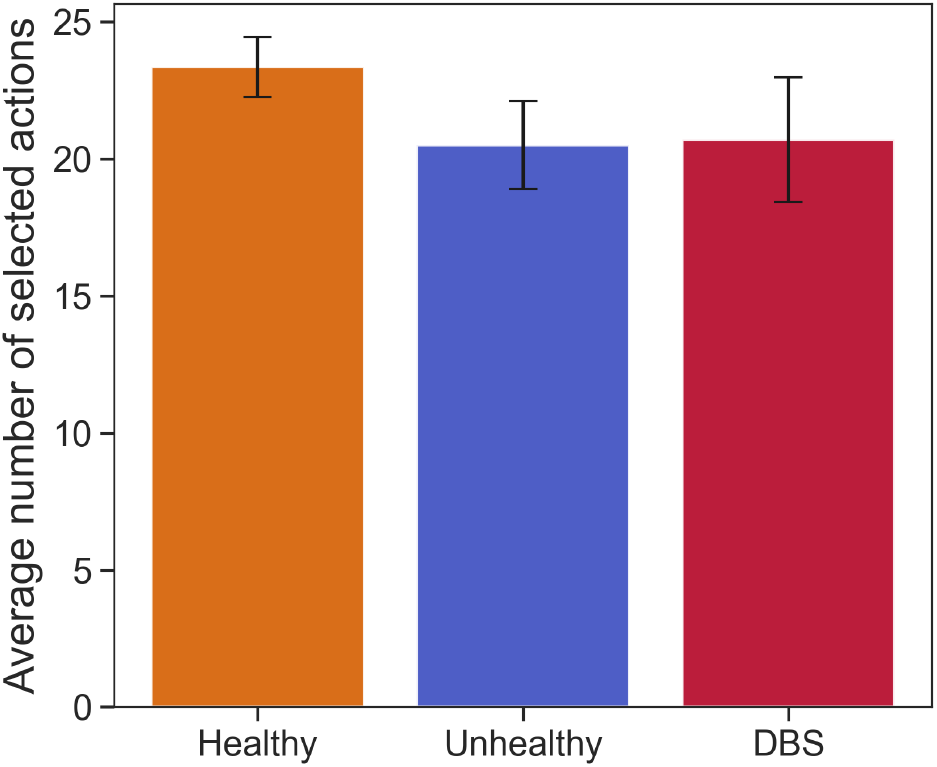
Mean number of actions selected (out of 25) under healthy, dopamine-depleted, and DBS-stimulated conditions for the representative parameter pair (*w* _*p*_, *w*_*t*_) = (0.70, 0.60). Bars show averages across *n* = 20 experiments, with error bars indicating ±1 standard deviation of action selection counts across repeated experiments for each condition.

## Discussion

This work investigated dopaminergic modulation and action selection dynamics in Bartlett et al. (2025)’s DNF-BG model, with two primary goals: (i) to validate whether the model robustly differentiates healthy and Parkinsonian conditions, and (ii) to explore whether DBS, modeled as modulation to inhibit the afferent connection to STN and GPi, can restore thalamic gating and successful action selection under Parkinsonian conditions to alleviate freezing symptoms.

Results from Study 1 demonstrate that the DNF-BG model reliably distinguishes healthy from dopamine-depleted conditions across a substantial parameter space. In particular, higher thalamic thresholds *θ* combined with high DNF scaling factor *α* produced stable selection behavior under healthy dopamine levels, while demonstrating suppressed selection in the Parkinsonian model (see Figure 2), which we argue demonstrates successful modeling of freezing occurrences. The statistical analysis of the representative configuration within Study 2 also indicated significant difference between the healthy and Parkinsonian conditions.

In contrast to the clear boundary observed in Study 1, DBS effects in Study 2 were weak and inconsistent across the explored parameter space. Although isolated experiments exhibited modest improvements in action selection success between Parkinsonian and DBS conditions, the DBS grid search did not reveal a coherent region of reliable recovery, and the statistical analysis indicated no significant difference between conditions.

Importantly, these findings should not be interpreted as evidence against the efficacy of DBS within the model, but since the DNF-BG model framework differs architecturally from the one used in Senft et al., 2018’s paper, there needs to be an exploration of alternative DBS formulations that could be more effective. In this exploratory study, DBS was approximated as a static scaling of afferent inputs to the STN and GPi, capturing gross excitatory or inhibitory effects, but there are many different theories of DBS action, ranging from effective lesion models to formulations that explicitly incorporate axonal activation and time-varying stimulation dynamics (Ashkan et al., 2017; McIntyre & Foutz, 2013). Electrical stimulation may excite or inhibit neurons, interfere with depolarization, or alter network-level dynamics in more complex ways (Agnesi et al., 2013; Chiken & Nambu, 2016; Herrington et al., 2016; McIntyre et al., 2004). Consequently, it is still debated whether DBS functions primarily by reducing overall activity or by reorganizing pathological firing patterns into more functional dynamics. Thus, the absence of robust recovery under this simplified formulation suggests that effective DBS for this framework may depend on more nuanced interactions with pathological basal ganglia dynamics than static gain reduction alone.

Recent computational work by Maith et al. (2025) argues that DBS should not be interpreted as a purely suppressive or lesion-like mechanism. Instead, DBS is proposed to operate as a dual-process intervention combining local somatic suppression with axonal activation that reorganizes information flow through the basal ganglia–thalamocortical loop. In their framework, somatic suppression attenuates pathological firing patterns, while concurrent activation of efferent and passing fibers preserves or enhances downstream signal transmission. Translating this perspective to the present model suggests a natural extension in which DBS is decomposed into two interacting components,

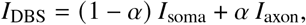

where *α* ∈ [0, 1] controls the balance between suppressive and excitatory neuron-mediated effects. Such a formulation would allow the parameters *w* _*p*_ and *w*_*t*_ to differentially influence somatic and axonal pathways, rather than uniformly scaling basal ganglia output.

Incorporating this dual-process view could offer a principled explanation for the limited improvements observed in the present DBS grid search. Selective modulation of inhibitory and excitatory gains captures only one dimension of DBS action, whereas stable recovery of action selection may depend on restoring effective communication across distributed pathways. Future work integrating explicit axonal activation mechanisms within the DNF-BG framework may therefore reveal broader regions of functional recovery, reduce trial-level variability, and clarify how DBS reshapes competition and convergence in continuous action selection. This is something to be explored in future work.

## Conclusion

In this work, we demonstrated that a Dynamic Neural Field–based basal ganglia model provides a robust computational account of Parkinsonian freezing as a failure of thalamic action gating under dopamine depletion. By coupling continuous action representations with an explicit thalamic gating network and threshold mechanism, the model reliably distinguished healthy and Parkinsonian regimes, reproducing stable action selection under intact dopaminergic modulation and impaired, freeze-prone dynamics when dopamine was reduced (Ackermann et al., 1997; Bartlett et al., 2025; Erro et al., 2014; Gurney et al., 2001a; Obeso et al., 2008). These results validate the DNF-BG framework as a principled substrate for studying dopamine-dependent action selection and selection failures associated with Parkinsonian freezing.

An implementation of DBS, based on established computational formulations by Senft et al. (2018), that models deep brain stimulation as controlled modulation of afferent signals to affected basal ganglia nuclei, did not yield consistent restoration of healthy-like action gating within the DNF-BG model. While sparse experiments were somewhat successful in improving action selection, DBS effects remained weak and non-significant despite our extended search for an optimal configuration. This outcome suggests that the adopted DBS formulation did not produce robust or consistent effects in the novel DNF-BG architecture. Perhaps, the effects may depend on more nuanced and complex interactions between inhibitory, excitatory, and axonal dynamics within the basal ganglia–thalamocortical loop (Agnesi et al., 2013; Chiken & Nambu, 2016; Herrington et al., 2016; McIntyre & Foutz, 2013).

Taken together, these findings establish a validated computational model of Parkinsonian freezing and provide a strong foundation for future work investigating more nuanced DBS mechanisms, offering a path toward deeper insight into how DBS interacts with pathological basal ganglia dynamics.

## Acknowledgments

This project was supported by collaborative research funding from the National Research Council of Canada’s Artificial Intelligence for Design program (AI4D-151-1).

## Notes

### Competing Interest Statement

The authors have declared no competing interest.

## References

Ackermann, H., Konczak, J., & Hertrich, I. (1997). The temporal control of repetitive articulatory movements in parkinson’s disease. Brain and Language, 56(2), 312–319. 10.1006/brln.1997.1851

Agnesi, F., Johnson, M. D., & Vitek, J. L. (2013). Deep brain stimulation: How does it work? In Handbook of clinical neurology (pp. 39–54, Vol. 116). Elsevier. 10.1016/B978-0-444-53497-2.00004-8

Ashkan, K., Rogers, P., Bergman, H., & Ughratdar, I. (2017). Insights into the mechanisms of deep brain stimulation. Nature Reviews Neurology, 13(9), 548–554. 10.1038/nrneurol.2017.105

Bartlett, M., Furlong, P. M., Stewart, T. C., & Orchard, J. (2025). A computational model of action specification in the basal ganglia. bioRxiv, 2025–08.

Bekolay, T., Bergstra, J., Hunsberger, E., Dewolf, T., Stewart, T. C., Rasmussen, D., Choo, X., Voelker, A. R., & Eliasmith, C. (2014). Nengo: A python tool for building large-scale functional brain models. Frontiers in Neuroinformatics, 7, 48. 10.3389/fninf.2013.00048

Chiken, S., & Nambu, A. (2016). Mechanism of deep brain stimulation: Inhibition, excitation, or disruption? The Neuroscientist, 22(3), 313–322. 10.1177/1073858415581986

Erro, R., Tedeschi, M. R., Vitale, C., Buonocore, S., & Orefice, G. (2014). A single-case rehabilitation program based on cueing for freezing of speech. European Journal of Physical and Rehabilitation Medicine, 50(5), 561–565. https://www.minervamedica.it/en/journals/europa-medicophysica/article.php?cod=R33Y2014N05A0561

Goetz, C. G., & Pal, G. (2014). Initial management of parkinson’s disease. BMJ, 349, g6258. 10.1136/bmj.g6258

Gurney, K., Prescott, T. J., & Redgrave, P. (2001a). A computational model of action selection in the basal ganglia. i. a new functional anatomy. Biological Cybernetics, 84, 401–410. 10.1007/PL00007984

Gurney, K., Prescott, T. J., & Redgrave, P. (2001b). A computational model of action selection in the basal ganglia. ii. analysis and simulation of behaviour. Biological Cybernetics, 84, 411–423. 10.1007/PL00007985

Herrington, T. M., Cheng, J. J., & Eskandar, E. N. (2016). Mechanisms of deep brain stimulation. Journal of Neurophysiology, 115(1), 19–38. 10.1152/jn.00281.2015

Humphries, M. D., Khamassi, M., & Gurney, K. (2012). Dopaminergic control of the exploration-exploitation trade-off via the basal ganglia. Frontiers in Neuroscience, 6, 16922.

Liu, Y., Li, W., Tan, C., Liu, X., Wang, X., Gui, Y., Qin, L., Deng, F., Hu, C., & Chen, L. (2014). Meta-analysis comparing deep brain stimulation of the globus pallidus and subthalamic nucleus to treat advanced parkinson disease. Journal of Neurosurgery, 121(3), 709–718. 10.3171/2014.4.JNS131711

Maith, O., Apenburg, D., & Hamker, F. H. (2025). Pallidal deep brain stimulation enhances habitual behavior in a neuro-computational basal ganglia model during a reward reversal learning task. European Journal of Neuroscience, 61(9), e70130. 10.1111/ejn.70130

McIntyre, C. C., & Foutz, T. J. (2013). Computational modeling of deep brain stimulation. Handbook of Clinical Neurology, 116, 55–61. 10.1016/B978-0-444-53497-2.00005-X

McIntyre, C. C., Savasta, M., Kerkerian-Le Goff, L., & Vitek, J. L. (2004). Uncovering the mechanism(s) of action of deep brain stimulation: Activation, inhibition, or both. Clinical Neurophysiology, 115(6), 1239–1248. 10.1016/j.clinph.2003.12.024

Mink, J. W. (1996). The basal ganglia: Focused selection and inhibition of competing motor programs. Progress in Neurobiology, 50(4), 381–425. 10.1016/S0301-0082(96)00042-1

Miocinovic, S., Parent, M., Butson, C. R., Hahn, P. J., Russo, G. S., Vitek, J. L., & McIntyre, C. C. (2006). Computational analysis of subthalamic nucleus and lenticular fasciculus activation during therapeutic deep brain stimulation. Journal of Neurophysiology, 96(3), 1569–1580. 10.1152/jn.00305.2006

Obeso, J. A., Rodríguez-Oroz, M. C., Benitez-Temino, B., Blesa, F. J., Guridi, J., Marin, C., & Rodriguez, M. (2008). Functional organization of the basal ganglia: Therapeutic implications for Parkinson’s disease. Movement Disorders, 23(Suppl 3), S548–S559. 10.1002/mds.22062

Redgrave, P., Prescott, T. J., & Gurney, K. (1999). The basal ganglia: A vertebrate solution to the selection problem? Neuroscience, 89(4), 1009–1023. 10.1016/S0306-4522(98)00319-4

Saha, P., Bartlett, M., & Jeff, O. (2026). A computational model of action selection and action specification in the basal ganglia.

Schöner, G., & Spencer, J. P. (2016). Dynamic thinking: A primer on dynamic field theory. Oxford University Press.

Senft, V., Stewart, T. C., Bekolay, T., Eliasmith, C., & Kröger, B. J. (2018). Inhibiting basal ganglia regions reduces syllable sequencing errors in parkinson’s disease: A computer simulation study. Frontiers in Computational Neuroscience, 12, 41. 10.3389/fncom.2018.00041

Vercruysse, S., Gilat, M., Shine, J. M., Heremans, E., Lewis, S. J. G., & Nieuwboer, A. (2014). Freezing beyond gait in parkinson’s disease: A review of current neurobehavioral evidence. Neuroscience & Biobehavioral Reviews, 43, 213–227. 10.1016/j.neubiorev.2014.04.010

Voelker, A. R., Blouw, P., Choo, X., Dumont, N. S.-Y., Stewart, T. C., & Eliasmith, C. (2021). Simulating and predicting dynamical systems with spatial semantic pointers. Neural Computation, 33(8), 2033–2067.

